# Non Parametric Differential Network Analysis: A Tool for Unveiling Specific Molecular Signatures

**DOI:** 10.1101/2024.04.29.591750

**Authors:** Pietro Hiram Guzzi, Roy Arkaprava, Marianna Milano, Pierangelo Veltri

## Abstract

The rewiring of molecular interactions in various conditions leads to distinct phenotypic outcomes. Differential Network Analysis (DNA) is dedicated to exploring these rewirings within gene and protein networks. Leveraging statistical learning and graph theory, DNA algorithms scrutinize alterations in interaction patterns derived from experimental data. Introducing a novel approach to differential network analysis, we incorporate differential gene expression based on sex and gender attributes. We hypothesize that gene expression can be accurately represented through non-Gaussian processes. Our methodology involves quantifying changes in non-parametric correlations among gene pairs and expression levels of individual genes. Applying our method to public expression datasets concerning diabetes mellitus and atherosclerosis in liver tissue, we identify gender-specific differential networks. Results underscore the biological relevance of our approach in uncovering meaningful molecular distinctions.

**Author summary:** This paper explores a novel technique for Differential Network Analysis (DNA) that considers sex-based variations. DNA compares biological networks under different conditions, like healthy vs. diseased states. Our method tackles the limitations of traditional DNA approaches, which often assume specific data distributions. We propose a non-parametric DNA methodology that integrates sex differences and identifies differential edges between networks. This approach utilizes data on gene expression levels and sex to construct a more accurate picture of the molecular mechanisms underlying diseases, particularly those exhibiting sex-dependent variations. Our method paves the way for a deeper understanding of how sex and age influence disease processes at the molecular level.

## Introduction

The emergence of high-throughput technologies in genomics, proteomics, and non-coding RNA studies has revolutionized our understanding of how variations in the abundance of these biological molecules correlate with diseases [1]. This deluge of data has spurred the creation of innovative analytical techniques that adopt a system-level approach through the lens of network science. These techniques employ networks to depict the complex interactions between biological molecules under specific conditions, as inferred from empirical data [2]. A particularly compelling use of network theory is comparing biological networks under disparate conditions, such as contrasting a disease state with a healthy one.

Differential Network Analysis (DNA) has gained recognition for its ability to delineate the differences between two states by encapsulating them within a singular differential network that highlights their variances [3–6]. DNA has found application in contrasting various experimental conditions or phenotypes, with recent studies underscoring the significance of factors like the age and sex of patients on drug response, disease progression, and comorbidity prevalence in chronic diseases [7–9]. Empirical findings have demonstrated notable disparities in disease incidence and progression based on sex and age, such as the heightened susceptibility of older diabetic patients to comorbidities and the observed sex-based differences in COVID-19 mortality rates [10–12]. This underscores the need for advanced algorithms to unravel the molecular mechanisms driving these age- and sex-dependent disparities.

DNA algorithms are designed to pinpoint changes in network structures by identifying association measures that differ between two biological states, *𝒞*_1_, *𝒞*_2_. When presented with two disparate biological conditions, represented by two networks of molecular interactions, DNA algorithms aim to uncover the network rewiring that underpins the mechanistic differences between these states. DNA algorithms construct networks *𝒩*_1_, *𝒩*_2_ for each condition, starting with gene expression datasets from two conditions. These networks feature nodes for each gene and weighted edges that denote the strength and nature of associations or causal relationships between genes. A differential network *𝒩*_*d*_ is then derived to represent the variation in associations across conditions, a technique previously applied to investigate disease-related alterations [8, 13].

Traditional methods often assume that gene expression data adhere to specific parametric distributions, such as Gaussian or Poisson distributions [1, 14–16]. However, the count-based nature of Next-Generation Sequencing (NGS) data challenges these assumptions, prompting the necessity for non-parametric DNA analysis approaches.

We introduce a cutting-edge DNA methodology that identifies differential edges between networks and integrates differential gene expressions, taking into account sex-based differences [12]. This approach employs multivariate count data for predicting gene expression levels and constructs a conditional dependence graph using pairwise Markov random fields [17]. This departure from traditional methods, which often presuppose parametric distributions for gene expression data, highlights the imperative for non-parametric techniques in DNA analysis.

Our proposed DNA algorithm is initiated by constructing two condition-specific graphs, from which a final differential graph is derived. This graph is then pruned to emphasize edges related to genes exhibiting differential expression. Our DNA method facilitates the identification of differential networks while incorporating considerations of gender differences, thereby advancing our understanding of the molecular basis of disease in the context of sex and age.

## Related Work

DNA’s application in distinguishing differentially expressed genes among various sample groups is invaluable, especially in contrasting individuals with specific diseases against healthy controls. This methodological approach is crucial in molecular biology and bioinformatics for pinpointing genes with variable expression levels between diseased and healthy sample groups.

Central to DNA-based research are algorithms designed to detect alterations in network structures under varying conditions [18]. These algorithms have been pivotal in biology for mapping the transition from healthy to diseased states within the same biological framework [2]. Our focus narrows to networks that maintain constant node sets yet exhibit variable edge sets. Specifically, in the presence of two distinct conditions *𝒞*_1_ and *𝒞*_2_, represented by graphs *G*_1_(*V, E*_1_) and *G*_1_(*V, E*_2_), the objective of DNA analysis is to pinpoint the modifications of the network.

In biological systems analysis, it is pertinent to note that while nodes represent directly quantifiable entities, the derivation of edges necessitates observing a sequence of temporal data. For instance, gene networks originating from microarray experiments necessitate the inference of edges from data through *statistical graphical models* [19–21]. In these models, each node within the graph *G* = (*V, E*) is one of these measurable random variables *X*_1_, …, *X*_*M*_, and the edges quantify a pre-specified notion of associations between the pairs of these variables. In this setting, the focus is predominantly on undirected graphs where the directions or the causality of these associations are not of interest. Among different metrics of associations, partial correlation is one of the most common ones as it measures conditional dependencies. Probabilistic graphical models allow conditional dependency-based graph estimation. Differential associations within these models are scrutinized by evaluating the variance in partial correlations across experimental conditions, utilizing specific statistical tests to measure the alterations in correlations among entities. Additionally, changes in gene expression levels are assessed using the classical Student’s t-test [22]. Subsequently, these statistical evaluations are amalgamated into a singular optimization model aiming to elucidate the hierarchical network structures. However, certain assumptions inherent to previous models, such as Gaussian data distribution, may not hold across all experimental conditions, necessitating non-parametric methods. While computationally efficient and simpler to implement, these methods demand adherence to specific distributional prerequisites, failing which could skew or invalidate the results.

Several studies have opted for a nonparanormal data distribution (or Gaussian copula) approach [23], employing rank-based correlation matrices like Spearman correlation or Kendall’s *τ*. There are other variations available too [18]. However, the nonparanormal models are primarily only suitable for continuous data, which limits its applicability in other settings.

These models have found applications in analyzing brain data and sequencing counts, circumventing the temporal limitations of non-parametric methods. Efficient Bayesian models have emerged [18], calculating edge probabilities by inferring their likelihood. Some methods adopt diverse heuristics for probability inference, challenging direct data derivation, as highlighted in [17], with this method surpassing other contemporary techniques.

Non-parametric methods, recognized for their minimal assumptions regarding data distribution, leverage data-driven approaches to evaluate network connectivity differences between conditions. Their flexibility and robustness are advantageous in handling complex, non-linear node relationships within networks, albeit at the cost of computational intensity and reduced interpretability.

The decision between parametric and non-parametric approaches for differential network analysis hinges on data characteristics, foundational assumptions, and the investigative query. Researchers frequently use sensitivity analysis and result cross-validation to ensure their findings’ robustness and reliability. Integrating insights from both methodological spectrums can yield a more detailed comprehension of the differential network architecture.

## Materials and methods

### Non Parametric Differential Network Analysis Algorithm

Let us consider two different expression datasets encoded in two matrices *N*_*j*_ *× M* (*N*_*j*_ samples, *M* genes) for *j* = 1, 2 denoted as *X*^1^, *X*^2^ representing two biological conditions *𝒞*_1_ *𝒞*_2_. Each row of *X*^*j*^ stores the expression values of *M* genes of different samples. Therefore, 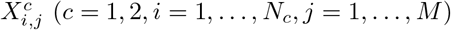 denotes expression of *j*-*th* gene in *i-th* sample under condition *c*. Note that the sample sizes under the two conditions may be different. We model this data under a Bayesian non-parametric framework.

Each column representing a gene may be encoded as a network node and compute conditional independence-based graphical relation. Let *M × M* dimensional matrices *𝒫*_1_ and *𝒫*_2_ represents the conditional independence relation among the *M* genes [17]. We then define the differential relation between two conditions based on the posterior samples of *𝒫*_1_ and *𝒫*_2_.

Following pairwise Markov random field (MRF) model for counts from [17], we consider the following joint probability mass function for *M*-dimensional count-valued data *X*,

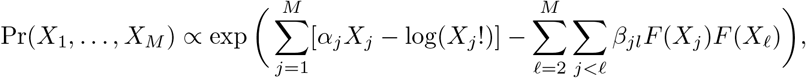

where *F* (*·*) is a monotone increasing bounded function with support [0, ∞). We let *F* (*·*) = (tan^−1^(*·*))^*θ*^ for some positive *θ* ∈ ℝ^+^ to define a flexible class of monotone increasing bounded functions. The exponent *θ* provides additional flexibility, including impacting the range of 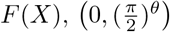. The parameter *θ* can be estimated along with the other parameters, including the baseline parameters *α* controlling the marginal count distributions and the coefficients *β*_*jl*_ controlling the graphical dependence structure. For simplicity, we pre-specify *θ* by estimating it as a minimizer of the loss, quantifying the difference in covariance between *F* (*X*) and *X* following [17]. For detailed descriptions of the method, readers are encouraged to check [17].

If *β*_*jℓ*_ = 0, we have *X*_*j*_ and *X*_*ℓ*_ to be conditionally independent, i.e. *P* (*X*_*j*_, *X*_*ℓ*_ | *X*_−(*j*,*ℓ*)_) = *P* (*X*_*j*_ | *X*_−(*j*,*ℓ*)_)*P* (*X*_*ℓ*_ | *X*_−(*j*,*ℓ*)_), where *X*_−(*j*,*ℓ*)_ stands for all the variables excluding *X*_*j*_ and *X*_*ℓ*_. Our estimated graphical relation would rely on *β*_*jℓ*_’s, and thus, our model falls under the probabilistic graphical model framework that encodes the conditional independence structure in a graph.

After running the above model under two conditions, we get the matrices *𝒫*_1_ and *𝒫*_2_ with (*j, k*)-*th* entries as 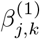 and 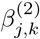 respectively. Consequently, a differential network is defined as the difference 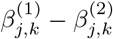 for each edge (*j, k*) where 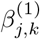 and 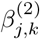 are estimated coefficients under two conditions 1 and 2. From the MCMC samples, we can get the posterior estimates of these differences as 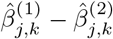. Alternatively, we can compute other posterior summaries such as 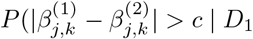, which is the posterior probability that 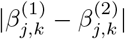 is greater than some pre-specified cutoff *c* given the two datasets, denoted as *D*_1_ and *D*_2_.

## Databases

The T2DiACoD database, as described by Rani et al. (2017) [24], was employed to collate a comprehensive list of genes linked to comorbid conditions associated with Type 2 Diabetes Mellitus (T2DM). Gene expression datasets were also sourced from the GTEx database [25].

T2DiACoD is a meticulously compiled database, the result of rigorous research and systematic literature review. It catalogues genes and noncoding RNAs that are crucial to understanding T2DM and its frequent comorbidities, including atherosclerosis, nephropathy, diabetic retinopathy, and cardiovascular disorders. This repository, enriched through meticulous data integration from existing databases, encapsulates 650 genes and 34 microRNAs related to these conditions, providing a reliable resource for your research.

The Genotype-Tissue Expression (GTEx) project is a vast open-access platform that empowers researchers like us with the distribution of genomic data collected from various individuals. This repository encompasses a broad spectrum of genomic data, from sequencing to methylation analyses, providing a wealth of information at your fingertips.

GTEx offers detailed metadata for each sample, covering aspects such as tissue type, sex, and age, categorized into six distinct groups. This makes GTEx an invaluable asset for investigating the interplay between age and tissue-specific gene expression. As of February 1st, the GTEx database boasts a collection of 17,382 samples across 54 tissue types from 948 donors, all accessible via the GTEx web portal. This portal enables users to efficiently search for and visualize data [12, 26, 27]. Furthermore, the data can be downloaded for in-depth analysis with custom scripts. In our research, we leveraged data from various tissues, dividing samples into two categories based on sex. Our study focused on the genes that play a pivotal role in developing T2DM-related complications, examining nine tissues (including blood, brain, adipose, amygdala, aorta, colon, coronary, liver, and lung tissues).

To ensure the validity of our research, we meticulously selected an equal number of samples from each tissue type, maintaining a balanced representation of age groups and uniformity in sample sizes across tissues. This approach facilitated an equitable distribution of age groups within each sex-based category for each tissue analyzed, ensuring the precision and reliability of our findings.

Focusing on atherosclerosis, we obtained a list of 115 genes related to this disease in T2DiACoD database, while for diabetes we obtained a list of 650 genes. We retrieved expression data by employing GTExVisualizer [8, 28], and metadata related to tissue, sex and age of the sample are extracted using genes identified in the T2DiACoD database in the previous step. Expression data are measured as Transcript per Million (TPM). This data integration and gene enrichment process was performed using an ad-hoc realised script that has been integrated into GTExVisualizer. We performed the analysis at tissue level, thus for each considered tissue, we split the data into male and female samples and randomly selected the same number of samples. We first generated DN by using non parametric methods and we evaluated the biological significanceby means of enrichment methods.

## Results

To show the effectiveness of our method we present two case studies on two chronic diseases to show differential mechanisms related to sex differences. We evaluated differential networks between men and women focusing on genes related to diabetes and atherosclerosis as reported in the T2DiaCoD database [24]. The characteristics of all the DNs are reported in Tab. 1 and 2. Then for each network, we performed a pathway enrichment analysis based on the KEGG pathway database [29] available on STRING enrichmennt app [30] of the Cytoscape software [31].

**Table 1.** Characteristics of the differential networks related to diabetes.

| DN | Nodes | Edges |
| --- | --- | --- |
| Liver | 128 | 3340 |
| Aorta | 237 | 2116 |
| Heart | 238 | 4316 |

**Table 2.** Characteristics of the differential networks related to atherosclerosis.

| DN | Nodes | Edges |
| --- | --- | --- |
| Adipose Visceral | 12 | 32 |
| Artery Coronary | 11 | 30 |
| Artery Tibial | 11 | 12 |
| Blood | 2 | 1 |

### Diabetes related Differential Networks

#### Liver Tissue

For liver tissue in diabetes, we obtained a DN with 128 nodes and 3340 edges, see Fig. 1. Fig. 16 depicts some subnetworks, the enrichment analysis highlighted the presence of some enriched pathways between sex.

**Fig 1.**
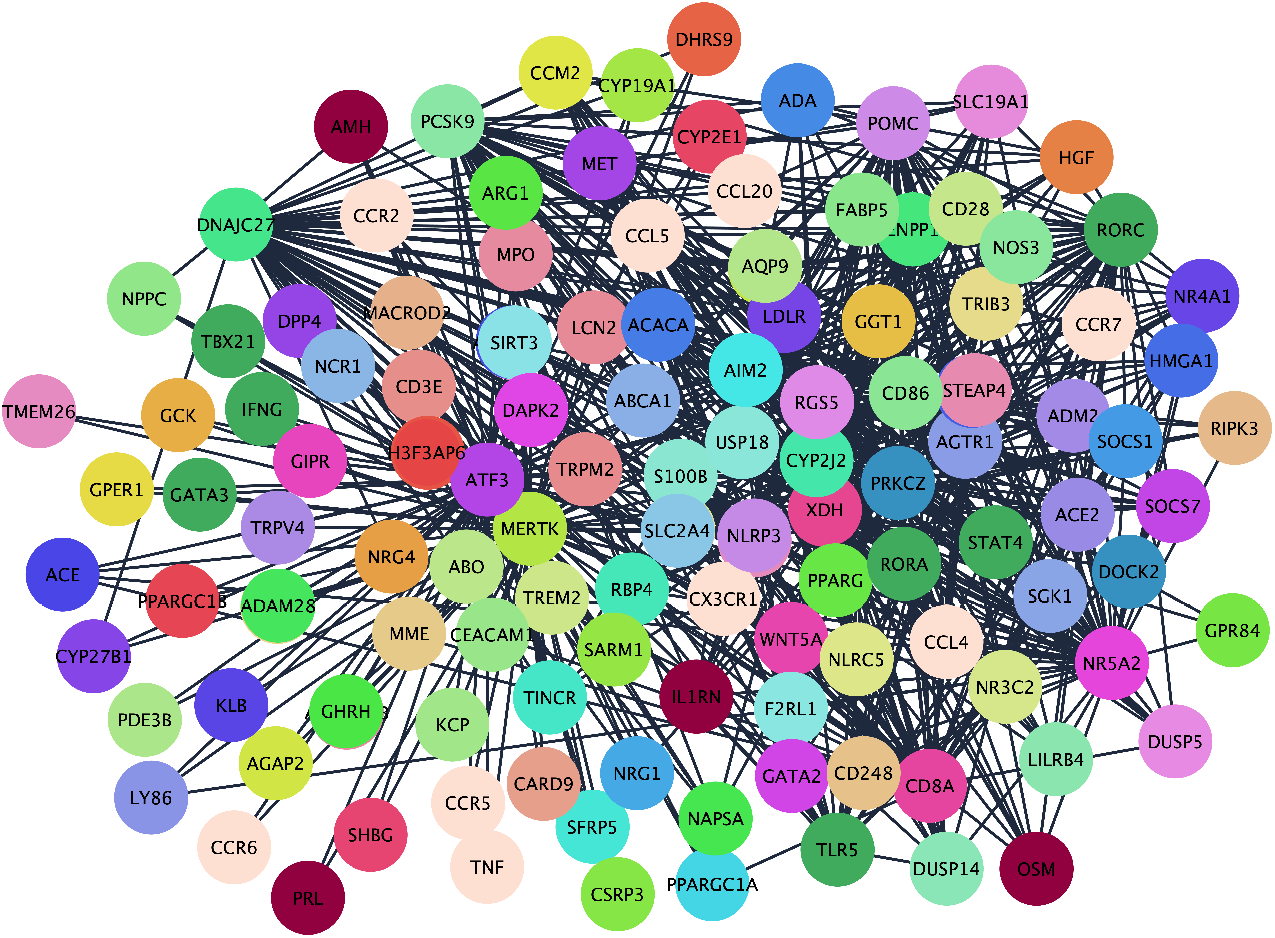
A select differential subnetwork for Liver tissue in Diabetes.

#### Aorta Tissue

For Aorta tissue in Diabetes we obtained a DN with 237 nodes and 2116 edges, see Fig. 2. Fig. 17 depicts some subnetworks, the enrichment analysis highlighted the presence of some enriched pathways between sex.

**Fig 2.**
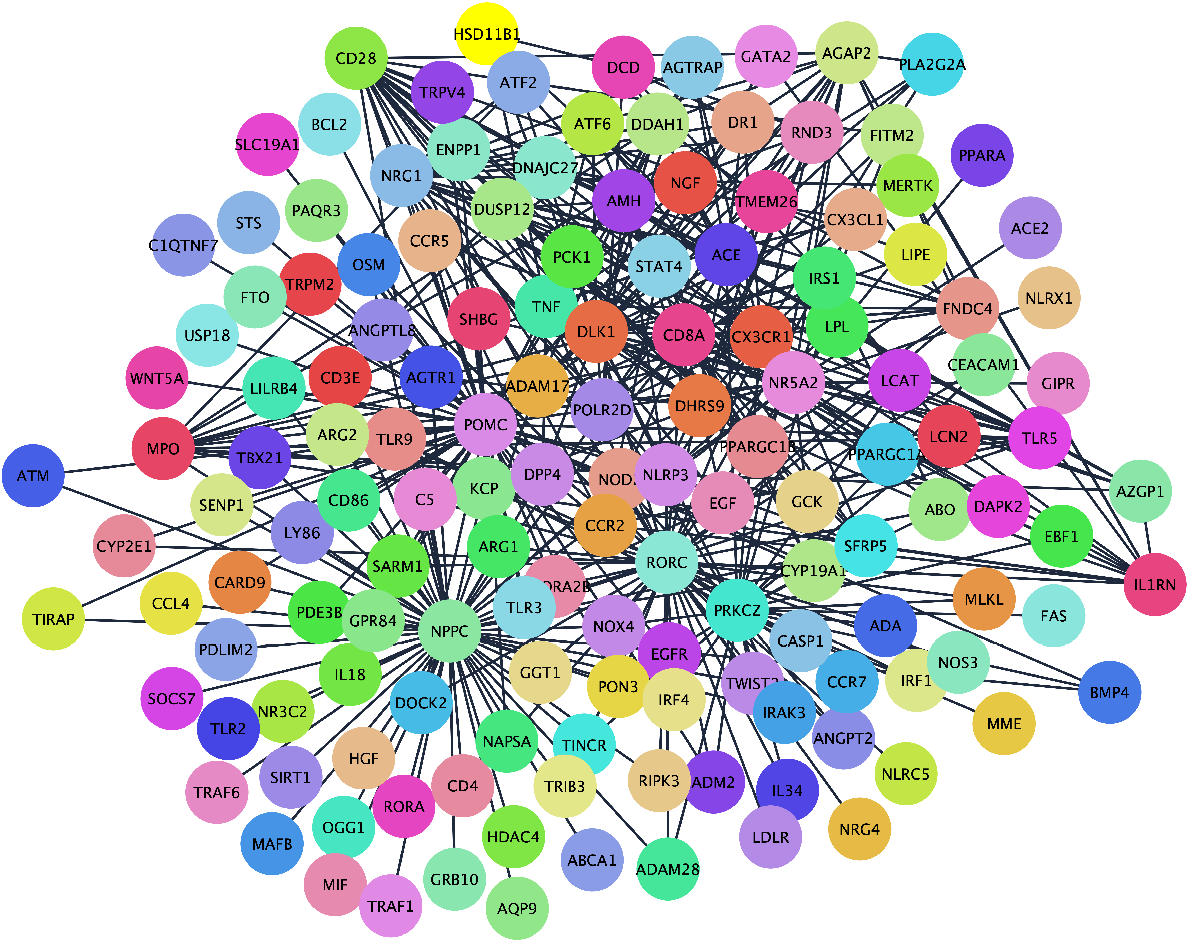
A select differential subnetwork for Aorta tissue in Diabetes.

#### Heart Tissue

For Hearth tissue in Diabetes we obtained a DN with 238 nodes and 4316 edges, see Fig. 3. Fig. 18 depicts some subnetworks, the enrichment analysis highlighted the presence of some enriched pathways between sex.

**Fig 3.**
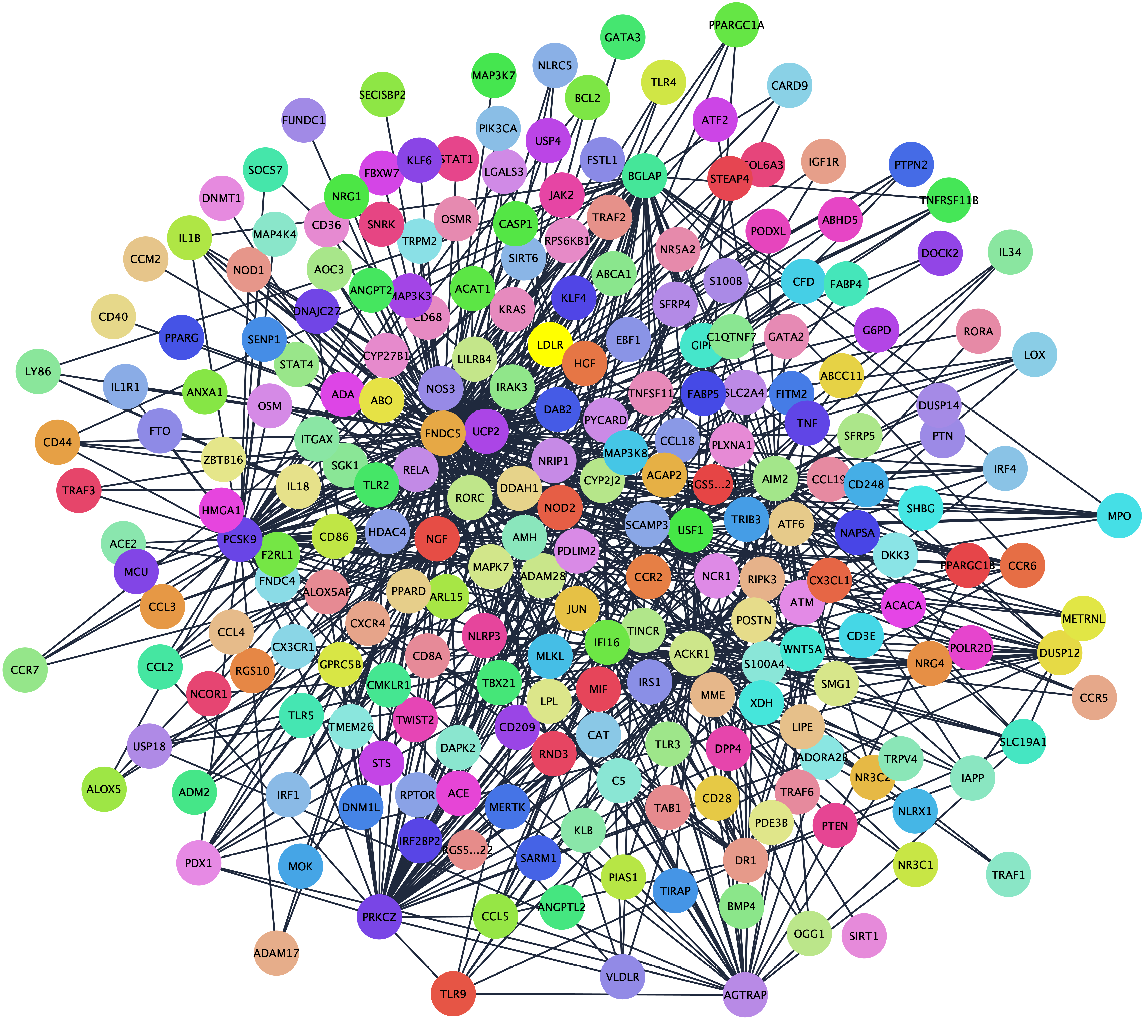
A select differential subnetwork for Heart tissue in Diabetes.

Fig. 8 shows a selected subnetwork of the differential network in Liver, Hearth and Aorta Tissue.

**Fig 4.**
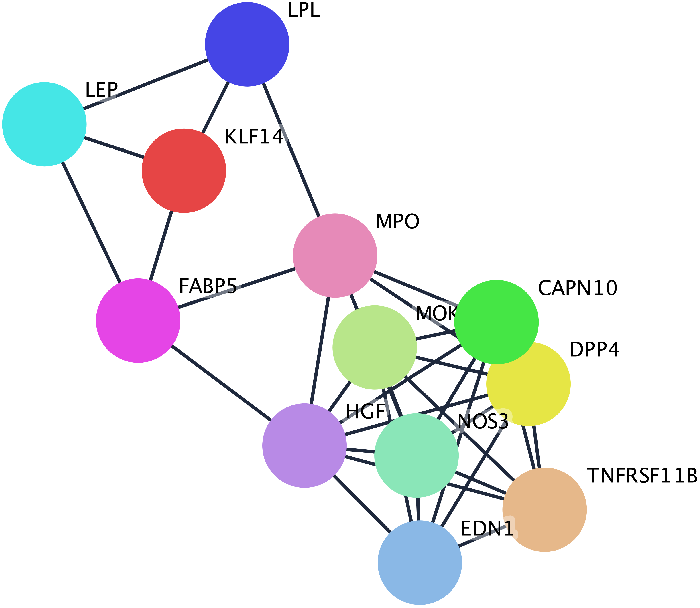
Differential network for Adipose Visceral tissue in Atherosclerosis.

**Fig 5.**
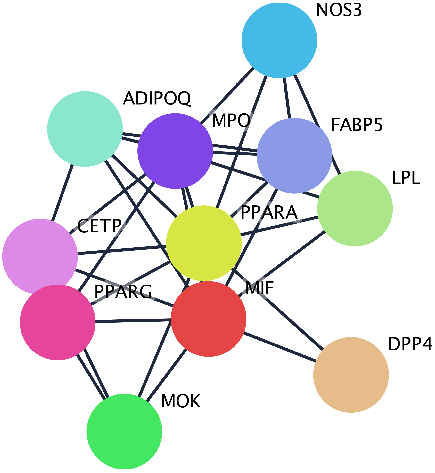
Differential network for Artery Coronary tissue in Atherosclerosis.

**Fig 6.**
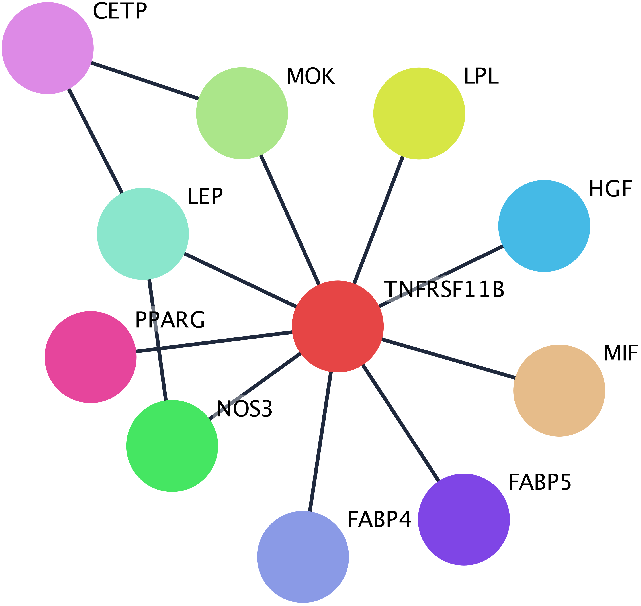
Differential network for Artery Tibial tissue in Atherosclerosis.

**Fig 7.**
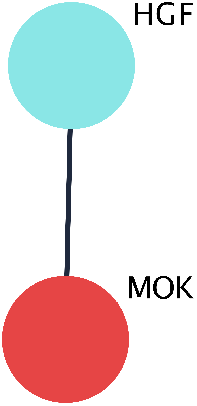
Differential network for Blood tissue in Atherosclerosis.

**Fig 8.**
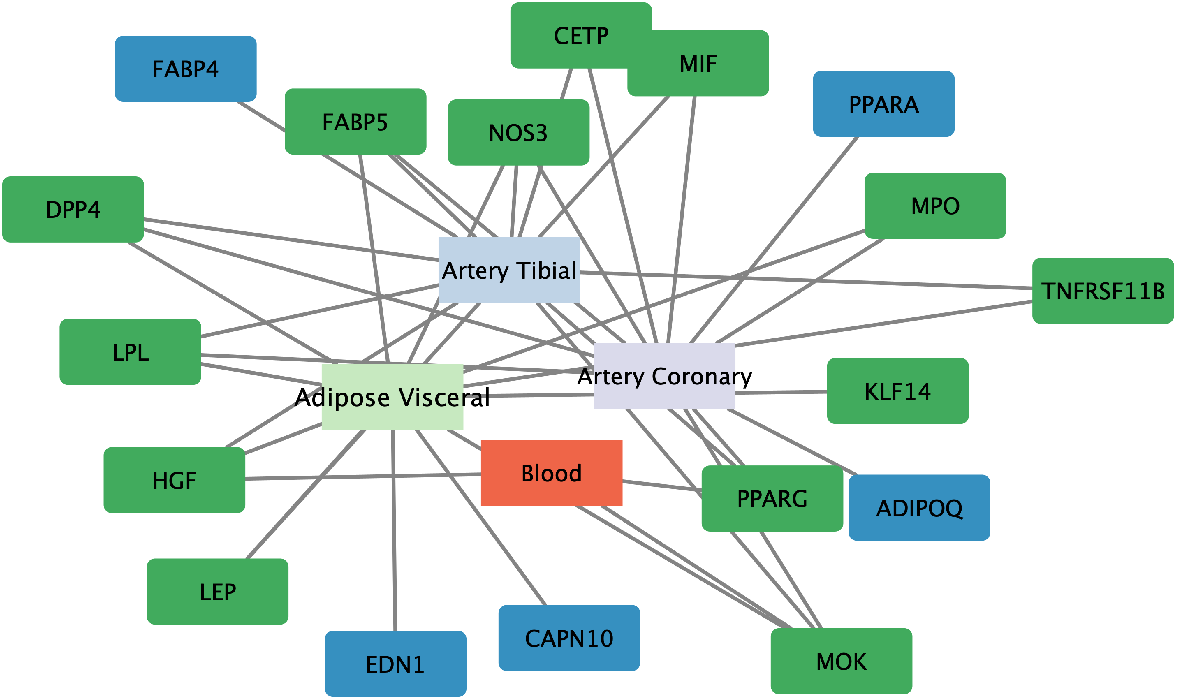
**Differential network related to diabetes expressed in Liver, Heart, Aorta tissues. The genes expressed in different tissues are reported in green, whereas the genes of specific tissue are reported in blue.**

Finally, we applied the Markov Clustering Algorithm to perform a cluster analysis on differential networks. Fig. 10, Fig. 11, Fig. 12 depict the clustered differential networks for Liver, Aorta, Heart tissues.

**Fig 9.**
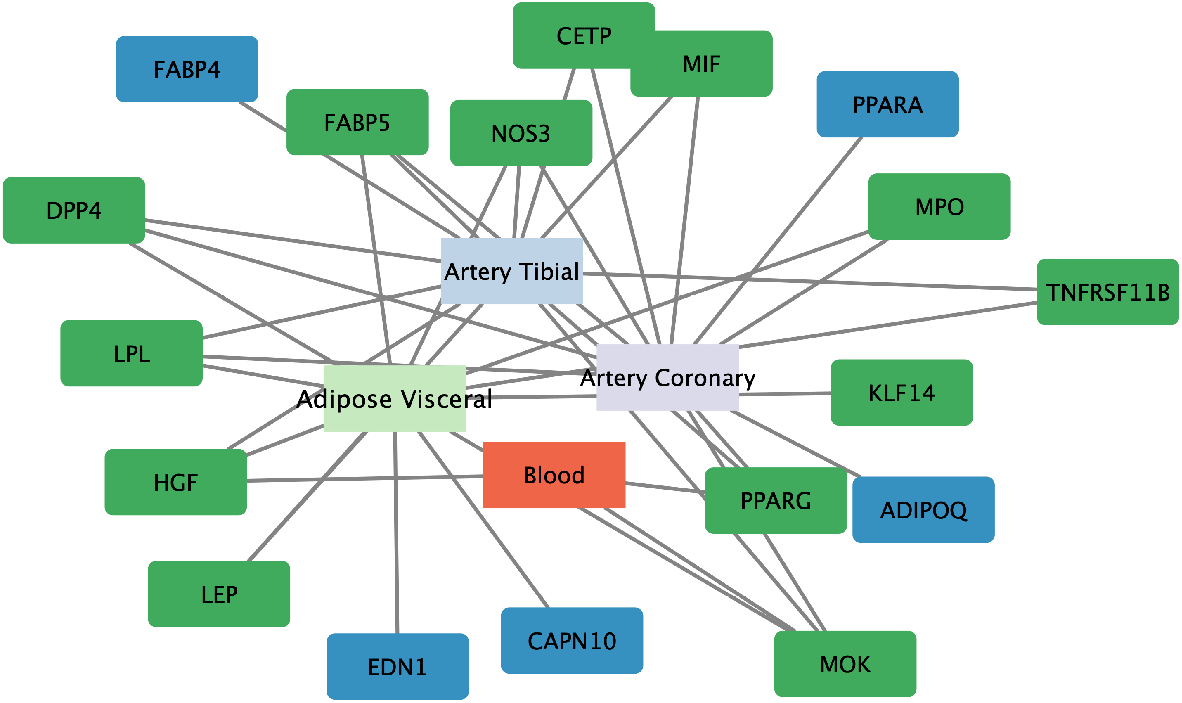
**Differential network related to atherosclerosis expressed in Adipose Visceral, Artery Coronary, Artery Tibial tissues. The genes expressed in different tissues are reported in green, whereas the genes of specific tissue are reported in blue.**

**Fig 10.**
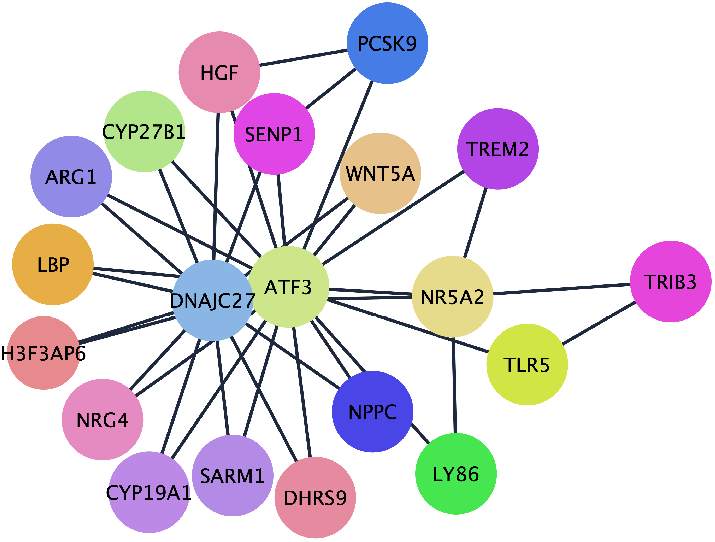
Clustered Differential subnetwork for Liver tissue in Diabetes.

**Fig 11.**
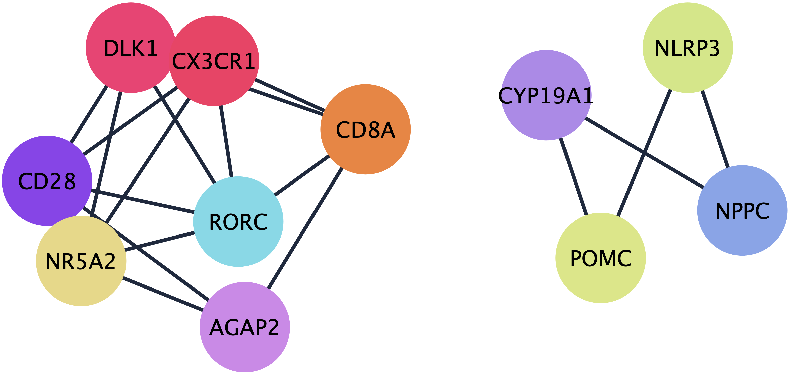
Clustered Differential subnetwork for Aorta tissue in Diabetes.

**Fig 12.**
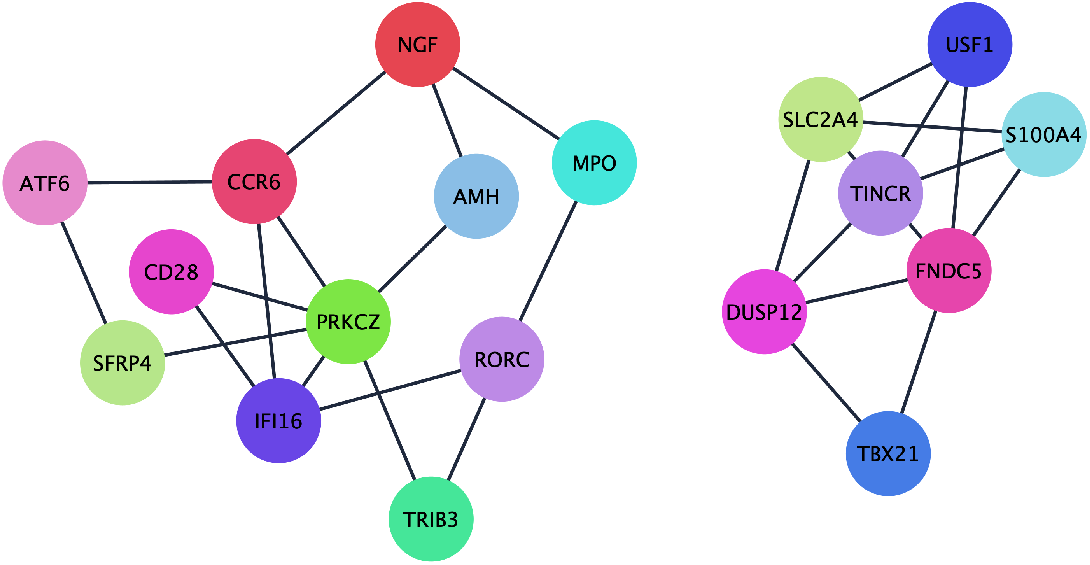
Clustered Differential subnetwork for Heart tissue in Diabetes.

### Atherosclerosis related Differential Networks

#### Adipose Visceral Tissue

For Adipose Visceral tissue in Atherosclerosis we obtained a DN with 12 nodes and 32 edges, see Fig. 4. The enrichment analysis did not report the presence of some enriched pathways between sex.

#### Artery Coronary Tissue

For Artery Coronary tissue in Atherosclerosis we identified a network of 11 nodes and 30 edges, see Fig. 5. Fig. 19 depicts some subnetworks, the enrichment analysis highlighted the presence of some enriched pathways between sex.

#### Artery Tibial Tissue

For Artery Tibial tissue in Atherosclerosis we identified a network of 11 nodes and 12 edges, see Fig. 6. Fig. 20 depicts some subnetworks, the enrichment analysis highlighted the presence of some enriched pathways between sex.

#### Blood Tissue

For Blood tissue in Atherosclerosis we identified a network of 2 nodes and 1 edges, see Fig. 7. The enrichment analysis did not report the presence of some enriched pathways between sex.

We summarize in Fig. 9 a selected subnetwork of the differential network in Adipose Visceral, Artery Coronary, and Aorta Tissue, Artery Tibial and Blood tissues.

Finally, we applied the Markov Clustering Algorithm to perform a cluster analysis on differential networks. Fig. 13, Fig. 14, Fig. 15 depict the clustered differential networks for Adipose Visceral, Artery Coronary, Artery Tibial tissues.

**Fig 13.**
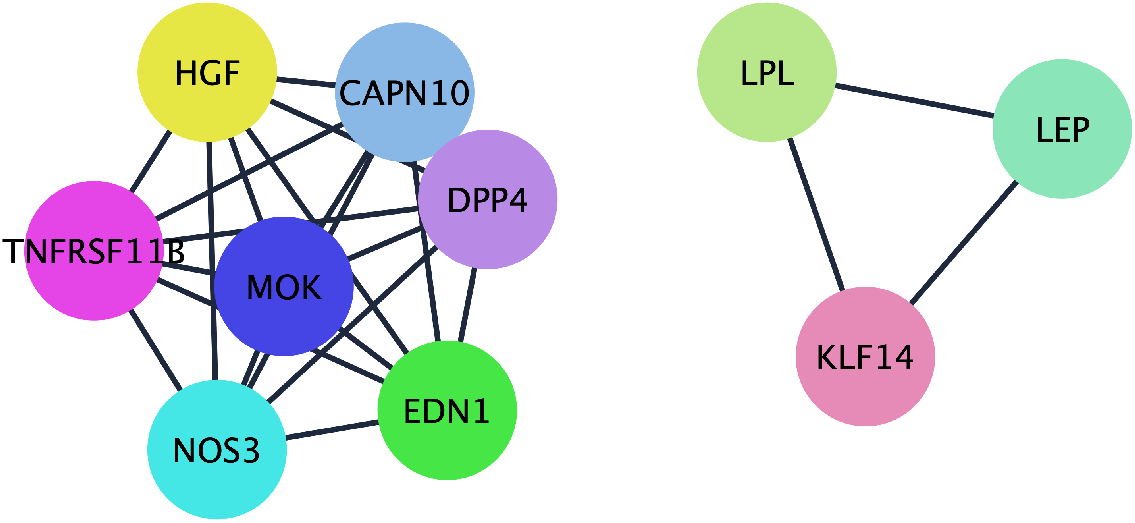
Clustered Differential network for Adipose Visceral tissue in Atherosclerosis.

**Fig 14.**
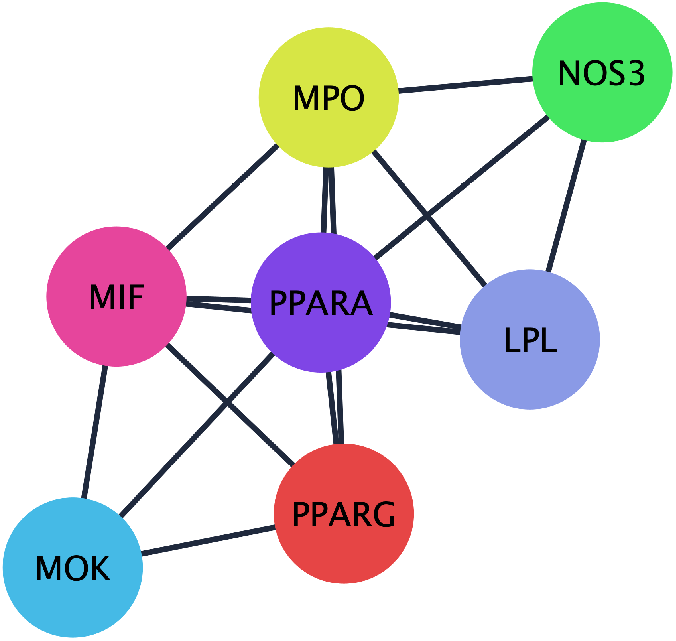
Clustered Differential network for Artery Coronary tissue in Atherosclerosis.

**Fig 15.**
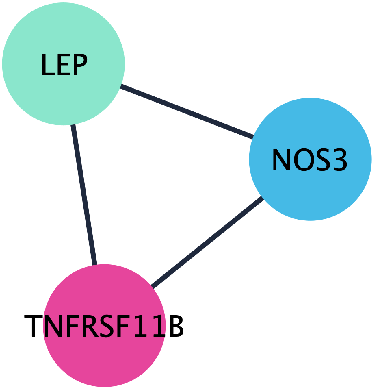
Clustered Differential network for Artery Tibial tissue in Atherosclerosis.

**Fig 16.**
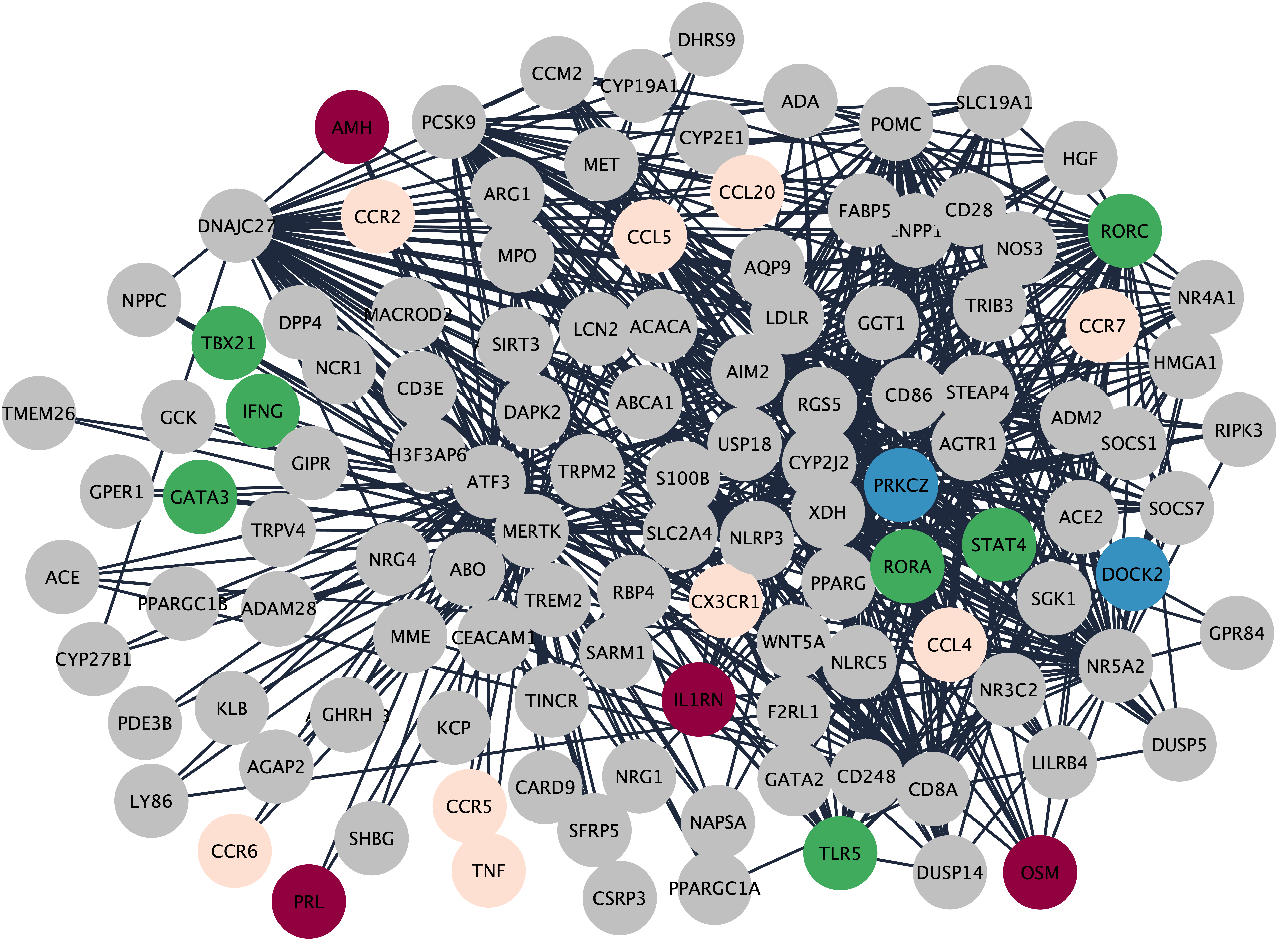
**A Selected subnetwork of the differential network for Liver tissue in Diabetes. We reported in different colour the genes involved in the pathways resulted from enriched analysis. Genes involved in Cytokine-cytokine receptor interaction are coloured in purple. Genes involved in Inflammatory bowel disease are coloured in green. Genes involved in Viral protein interaction with cytokine and cytokine receptor are coloured in pink. Genes involved in Chemokine signaling pathway are coloured in blue.**

**Fig 17.**
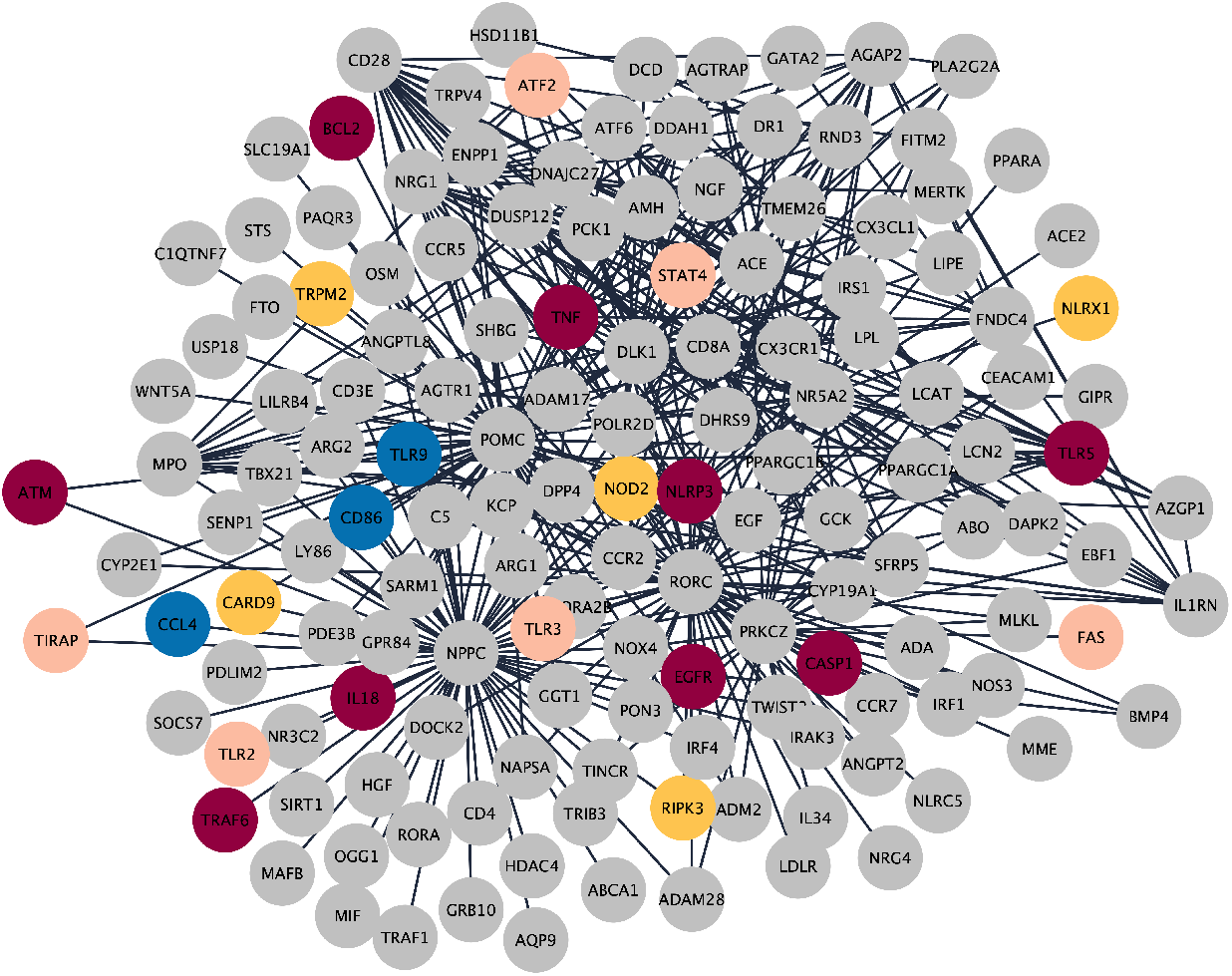
**A Selected subnetwork of the differential network for Aorta tissue in Diabetes. We reported in different colour the genes involved in the pathways resulted from enriched analysis. Genes involved Toll-like receptor signaling pathway are coloured in blue. Genes involved in NOD-like receptor signaling pathway are coloured in yellow. Genes involved in Hepatitis B are coloured in pink. Genes involved in Shigellosis are coloured in purple.**

**Fig 18.**
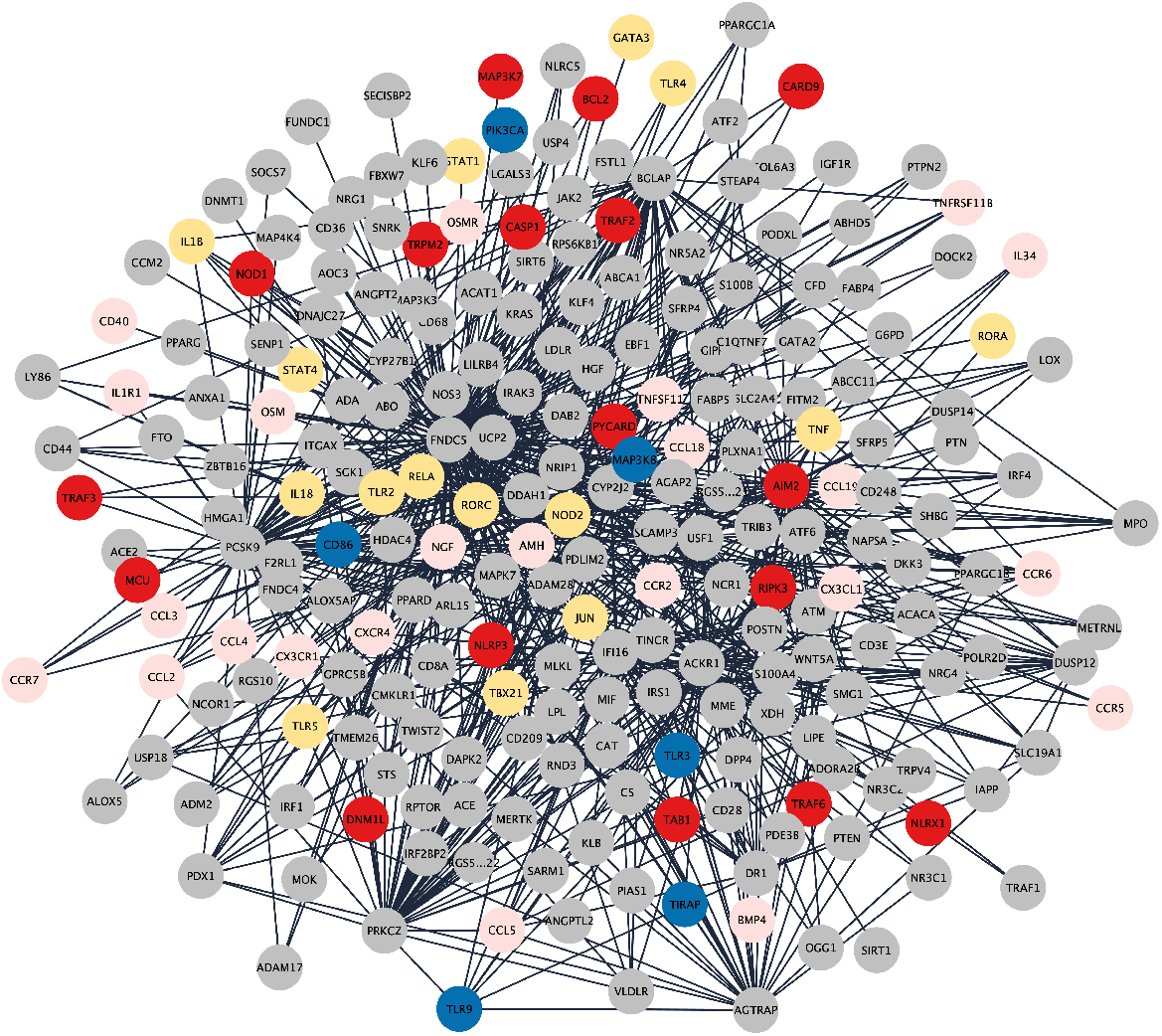
**A Selected subnetwork of the differential network for Heart tissue in Diabetes. We reported in different colour the genes involved in the pathways resulted from enriched analysis. Genes involved Toll-like receptor signaling pathway are coloured in blue. Genes involved in NOD-like receptor signaling pathway are coloured in red. Genes involved in Cytokine-cytokine receptor interaction are coloured in pink. Genes involved in Inflammatory bowel disease are coloured in yellow.**

**Fig 19.**
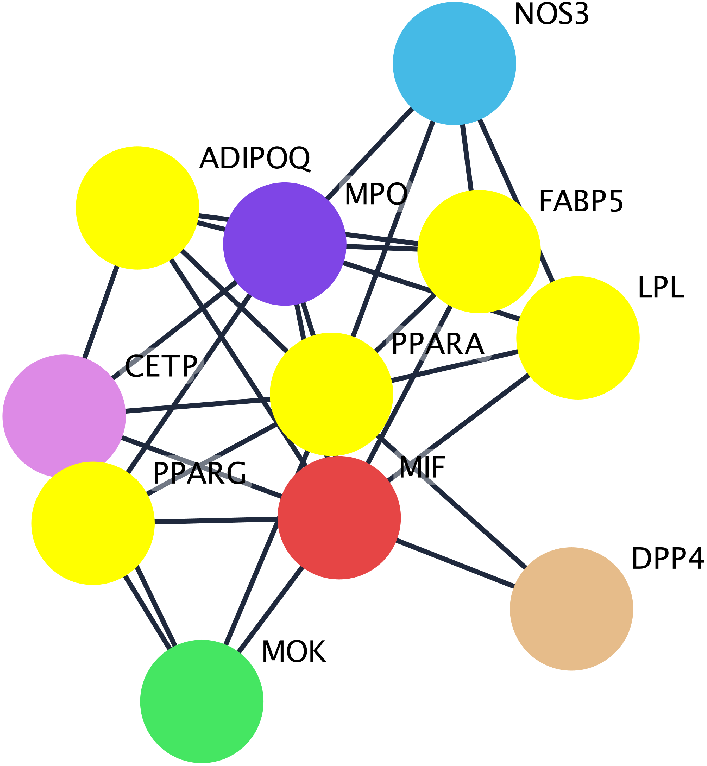
**A Selected subnetwork of the differential network for Artery Coronary tissue in Atherosclerosis. We reported in different colour the genes involved in the pathways resulted from enriched analysis. Genes involved in PPAR signaling pathway are reported in yellow.**

**Fig 20.**
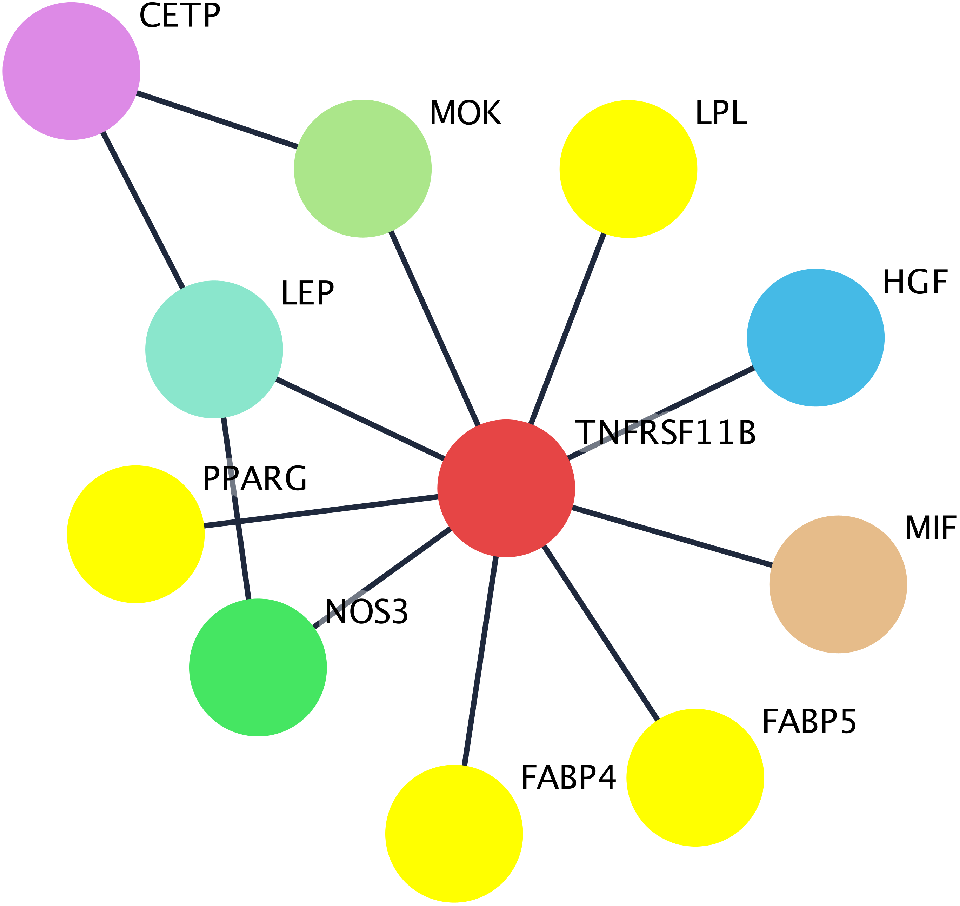
**A Selected subnetwork of the differential network for Artery Tibial tissue in Atherosclerosis. We reported in different colour the genes involved in the pathways resulted from enriched analysis. Genes involved in PPAR-***γ* **signaling pathway are reported in yellow.**

## Discussion

In our research study, we introduced a groundbreaking method called differential network analysis. This innovative approach is particularly beneficial for datasets that contain count data or non-parametric data, which are prevalent in biological research but pose challenges due to their non-normal distribution and discrete nature. By employing differential network analysis, we can delve into the underlying network structures between different conditions, providing a more comprehensive understanding of biological variations and interactions that are not discernible through traditional methods. This pioneering methodology is the first significant contribution of our paper, setting the stage for subsequent analyses and discussions.

Given the absence of an established standard in this area, the rigorous evaluation of the biological relevance of the networks derived from our differential network analysis was paramount. We meticulously scrutinized the networks’ components and connections, ensuring their biological significance and consistency with known biological pathways and mechanisms. This comprehensive analysis not only validated the practical utility of our methodology but also provided novel insights into the biological systems under study. By correlating our findings with existing biological knowledge and experimental data, we solidified the biological relevance of our results, further underscoring the value of differential network analysis in uncovering meaningful biological patterns and interactions. In the context of diabetes, our research yielded significant findings. We discovered that the differential networks in Liver, Aorta, and Heart tissues are enriched for the Toll-like receptor signaling pathway, as indicated in Tab. 3, Tab. 4, Tab. 5. The toll-like receptor 4 (TLR-4) pathway has been associated with various pathophysiological conditions, including cardiovascular diseases (CVDs) and Rheumatoid Arthritis (RA), underscoring the relevance of our findings in the broader context of human health. Different studies demonstrated that TLR4 activates the expression of several pro-inflammatory cytokine genes that play pivotal roles in myocardial inflammation, particularly myocarditis, myocardial infarction, ischemia-reperfusion injury, and heart failure [32–35].

Also, we found in Aorta and Heart tissues the NOD-like receptor signalling pathway; see Tab. 4 and Tab. 5. The NOD-like Receptor (NLR) family of proteins is a group of pattern recognition receptors (PRRs) known to mediate the initial innate immune response to cellular injury and stress. Different studies reveal the role of the activation of the Nod-like receptor protein 3 (NLRP3) inflammasome in the pathogenesis of many metabolic diseases, including diabetes and its complications [36].

Furthermore, the differential networks in the Aorta and Heart tissues were enriched for the TNF signalling pathway; see Tab. 4 and Tab. 5. Lamki et al. [37] reported that Tumor necrosis factor (TNF) represent a central mediator of a broad range of biological activities from cell proliferation, cell death and differentiation to induction of inflammation and immune modulation. TNF mediates the inflammatory response and regulates immune function. Inappropriate production of TNF or sustained activation of TNF signalling has been implicated in the pathogenesis of a broad spectrum of human diseases, including diabetes, cancer, osteoporosis, allograft rejection, and autoimmune diseases such as multiple sclerosis, rheumatoid arthritis, and inflammatory bowel diseases [37], [38].

**Table 3.** Top Enriched Pathways of liver tissue in Diabetes.

| Description | Term name |
| --- | --- |
| Cytokine-cytokine receptor interaction | hsa04060 |
| Viral protein interaction with cytokine and cytokine receptor | hsa04061 |
| Chemokine signaling pathway | hsa04062 |
| Inflammatory bowel disease | hsa05321 |
| Toll-like receptor signaling pathway | hsa04620 |
| Malaria | hsa05144 |
| Rheumatoid arthritis | hsa05323 |
| Renin-angiotensin system | hsa04614 |
| Th17 cell differentiation | hsa04659 |
| Chagas disease | hsa05142 |

**Table 4.** Top Enriched Pathways of aorta tissue in Diabetes.

| Description | Term name |
| --- | --- |
| Toll-like receptor signaling pathway | hsa04620 |
| NOD-like receptor signaling pathway | hsa04621 |
| Shigellosis | hsa05131 |
| Hepatitis B | hsa05161 |
| Salmonella infection | hsa05132 |
| TNF signaling pathway | hsa04668 |
| NF-kappa B signaling pathway | hsa04064 |
| Toxoplasmosis | hsa05145 |
| PD-L1 expression and PD-1 checkpoint pathway in cancer | hsa05235 |
| PI3K-Akt signaling pathway | hsa04151 |

**Table 5.** Top Enriched Pathways of heart tissue.

| Description | Term name |
| --- | --- |
| NOD-like receptor signaling pathway | hsa04621 |
| Toll-like receptor signaling pathway | hsa04620 |
| TNF signaling pathway | hsa04668 |
| Cytokine-cytokine receptor interaction | hsa04060 |
| NF-kappa B signaling pathway | hsa04064 |
| Salmonella infection | hsa05132 |
| Inflammatory bowel disease | hsa05321 |
| Shigellosis | hsa05131 |
| PD-L1 expression and PD-1 checkpoint pathway in cancer | hsa05235 |
| Viral protein interaction with cytokine and cytokine receptor | hsa04061 |

Instead, for the differential networks in Liver tissue were reported the chemokine signalling pathway that promotes changes in cellular morphology [39] and insulin signalling pathway that accounts for selective insulin resistance [40], see Tab. 4.

Also, the differential network in the Aorta tissue was enriched for the PI3K-Akt signalling pathway, see Tab. 5. Phosphatidylinositol 3-kinases (PI3Ks) are crucial coordinators of intracellular signalling in response to extracellular stimulators. The hyperactivation of PI3K signalling cascades is one of the most common events in human cancers. The high recurrence of phosphoinositide 3-kinase (PI3K) pathway adjustments in cancer has led to a surge in the progression of PI3K inhibitors [41]. For diabetes, we found that differential networks in Artery Coronary and Artery Tibial tissue were enriched for the PPAR signalling pathway, which activation is linked to a correlation between metabolic syndromes and cancer, see Tab. 6.

**Table 6.** Enriched Pathway of Artery Coronary and Artery Tibial tissue.

| Description | Term name |
| --- | --- |
| PPAR signaling pathway | hsa03320 |

## Conclusion

Differential Network Analysis (DNA) is a crucial tool for understanding the intricate interactions within biological systems, especially regarding specific conditions or phenotypes. This study utilizes DNA to explore the differential molecular interactions related to diabetes mellitus, taking into account age and gender-specific variations. Differential networks are critical since they enable the detection and visualisation of variations in gene and protein interactions between males and females. This innovative approach is essential for identifying subtle biological discrepancies that conventional analytical methods could miss. Non-parametric methods for DNA are emphasized in the study since they enhance the robustness and applicability of the findings, given the complex nature of biological data, rather than assuming normal distributions of gene expression data.

The biological significance of the DNA findings is significant, particularly with the integration of age and sex factors into the analysis. The study applies this method to liver tissue gene expression data related to diabetes, identifying distinct networks that may be involved in gender-specific disease mechanisms. These findings can provide insight into why certain diseases exhibit different manifestations in males and females, ultimately leading to more targeted approaches in treatment and management. Identifying gender-specific differential networks aids in understanding the molecular basis of diabetes and its variation with sex, providing potential pathways for therapeutic intervention and a deeper understanding of the disease’s etiology.

This study demonstrates the effectiveness of the differential network approach in discriminating between male and female biological samples. The method effectively identifies and visualizes differences in molecular interactions related to diabetes in liver tissue between the two sexes. Such discrimination is vital for personalized medicine, leading to more precise diagnostic and therapeutic strategies. By effectively highlighting the differences in gene expression and interactions, the study supports the potential of DNA in improving our understanding of sex-specific traits in diseases, which is crucial for advancing gender-specific medicine.

The article presents a compelling argument for using differential networks in biological research, especially in complex diseases such as diabetes. Integrating non-parametric methods improves the analysis by accounting for the inherent complexities and variations in biological data often overlooked in parametric approaches. This research contributes significantly to our understanding of diabetes and sets a precedent for future studies exploring other complex diseases with potential variations in expression and interaction patterns across different groups.

## Acknowledgments

This work was funded by the Next Generation EU - Italian NRRP, Mission 4, Component 2, Investment 1.5, call for the creation and strengthening of ‘Innovation Ecosystems’, building ‘Territorial R&D Leaders’ (Directorial Decree n. 2021/3277) - project Tech4You - Technologies for climate change adaptation and quality of life improvement, n. ECS0000009. This work reflects only the authors’ views and opinions, neither the Ministry for University and Research nor the European Commission can be considered responsible for them.

